# Exocytotic dynamics of glucagon-like peptide-1 from enteroendocrine L cell line is regulated by actin polymerization

**DOI:** 10.1101/2022.09.14.508035

**Authors:** Kazuki Harada, Maoko Takashima, Tetsuya Kitaguchi, Takashi Tsuboi

## Abstract

Stimulus-secretion coupling of glucagon-like peptide-1 (GLP-1) from enteroendocrine L cells is important for glucose homeostasis. Although intracellular second messengers including Ca^2+^ and cAMP, and cellular structures including actin cytoskeleton play roles in induction of exocytosis of GLP-1 granules, little is known about the specific part in the process of exocytosis in which they are involved. Here we explored the role of those molecules by live-cell imaging with mouse L cell line GLUTag cells, and used two stimuli: deoxycholic acid (DCA) and high K^+^. DCA increased both intracellular Ca^2+^ and cAMP levels, while high K^+^ only increased Ca^2+^. We next monitored a single exocytosis of GLP-1 granules and found that, during the first 10 minutes of stimulation, both stimuli mainly induced the exocytosis from the predocked granules with the plasma membrane before stimulation or granules immediately fused to the plasma membrane without docking. Furthermore, inhibition of actin polymerization suppressed the proportion of exocytosis by the predocked granules. These results suggest that the exocytotic process of GLP-1 granules is determined by interaction with F-actin upon the increase of either Ca^2+^ or cAMP.

**Summary statement:** Exocytotic process of glucagon-like peptide-1 granules from a mouse enteroendocrine L cell line is regulated by actin polymerization immediately after elevation of intracellular Ca^2+^ or cAMP levels.

## Introduction

Glucagon-like peptide-1 (GLP-1) is a peptide hormone produced in enteroendocrine L cells (Holst, 2007). GLP-1 secreted from L cells not only strengthens insulin secretion from pancreatic β cells but also induces stomach swelling and subsequent appetite suppression trough autonomous nervous system (Holst, 2007; Zhang et al., 2022). Furthermore, GLP-1 is produced in neurons in certain parts of the brain including the nucleus tractus solitarii, and acts as a neurotransmitter to exert anorexigenic effects (Trapp and Brierley, 2022). Because these functions of GLP-1 are helpful for the treatment of type 2 diabetes, diverse drugs such as dipeptidyl peptidase-4 inhibitor, GLP-1 receptor agonist, GLP-1 and glucose-dependent insulinotropic polypeptide (GIP) receptor dual agonist, and GLP-1, GIP and glucagon receptor triple agonist are in clinical use or trials (Bossart et al., 2022; Kalra et al., 2022; Villhauer et al., 2003).

GLP-1 secretion from L cells is achieved by exocytosis of GLP-1-containing granules. During exocytosis of GLP-1, granules in the cytoplasm are first recruited and bound to the plasma membrane (docking). Then, interaction of soluble NSF attachment protein receptor (SNARE) proteins transforms granules to a fusion-competent state (priming). Finally, granule membrane is fused with the plasma membrane and enables release of intra-granuler contents (fusion) (Seino and Shibasaki, 2005). Intracellular second messenger molecules including Ca^2+^ and cAMP, and cytoskeletons are major regulator of exocytosis (Miklavc and Frick, 2020; Shibasaki et al., 2007; Stožer et al., 2021). Studies on pancreatic β cells have revealed contribution of these molecules in spatial and temporal coordination of insulin exocytosis (Ma et al., 2020; Shibasaki et al., 2007; Yasuda et al., 2010; Yuan et al., 2015). Although there are several studies showing the importance of SNARE proteins and actin in the regulation of GLP-1 exocytosis (Campbell et al., 2020; Harada et al., 2018; Li et al., 2014; Wheeler et al., 2017), precise regulatory mechanism underlying exocytotic process of of GLP-1-containing granules is unknown.

In this study we aimed to elucidate the spatial and temporal regulation mechanism of GLP-1 exocytosis by live-cell imaging with mouse L cell line GLUTag cells. We used two kinds of stimuli: deoxycholic acid (DCA) and high K^+^. DCA is one of bile acids produced by gut microbiota, and known to induce GLP-1 secretion (Parker et al., 2012), while high K^+^ is generally used as control stimulation for excitatory cells including L cells (Psichas et al., 2017). DCA increased intracellular Ca^2+^ and cAMP levels ([Ca^2+^]_i_ and [cAMP]_i_, respectively) and high K^+^ increased [Ca^2+^]_i_ only. When we monitored a single GLP-1 exocytosis, both stimuli induced a similar spatial and temporal pattern of exocytosis types. Immediately after stimulation, released granules were mainly predocked to the plasma membrane or rapidly fused to the plasma membrane without docking. Moreover, inhibition of actin polymerization altered the observed patterns. We propose that cortical actin network plays a primary role in regulating the dynamics of GLP-1 exocytosis together with [Ca^2+^]_i_ and [cAMP]_i_.

## Results and Discussion

### Effect of DCA and high K^+^ on [Ca^2+^]_i_ and [cAMP]_i_

In order to distinguish between the role of Ca^2+^ and cAMP in the regulation of GLP-1 exocytosis, we chose DCA and high K^+^. We performed live-cell imaging to monitor intracellular Ca^2+^ dynamics with Ca^2+^-sensitive dye Fluo4, and cAMP dynamics with genetically-encoded cAMP indicator Pink Flamindo (Harada et al., 2017a), respectively. While 30 μM DCA caused an increase in the fluorescence intensity (FI) of both Fluo4 and Pink Flamindo (Fig. 1A to D), high K^+^ (15.2 mM KCl) only increased the FI of Fluo4 (Fig. 1E to H). Both DCA and high K^+^ increased the secretion of GLP-1 (Fig. 1I and J). These results suggest that DCA increases [Ca^2+^]_i_ and [cAMP]_i_, and high K^+^ only increases [Ca^2+^]_i_ in GLUTag cells, both resulting in enhancement of GLP-1 secretion.

**Figure 1.**
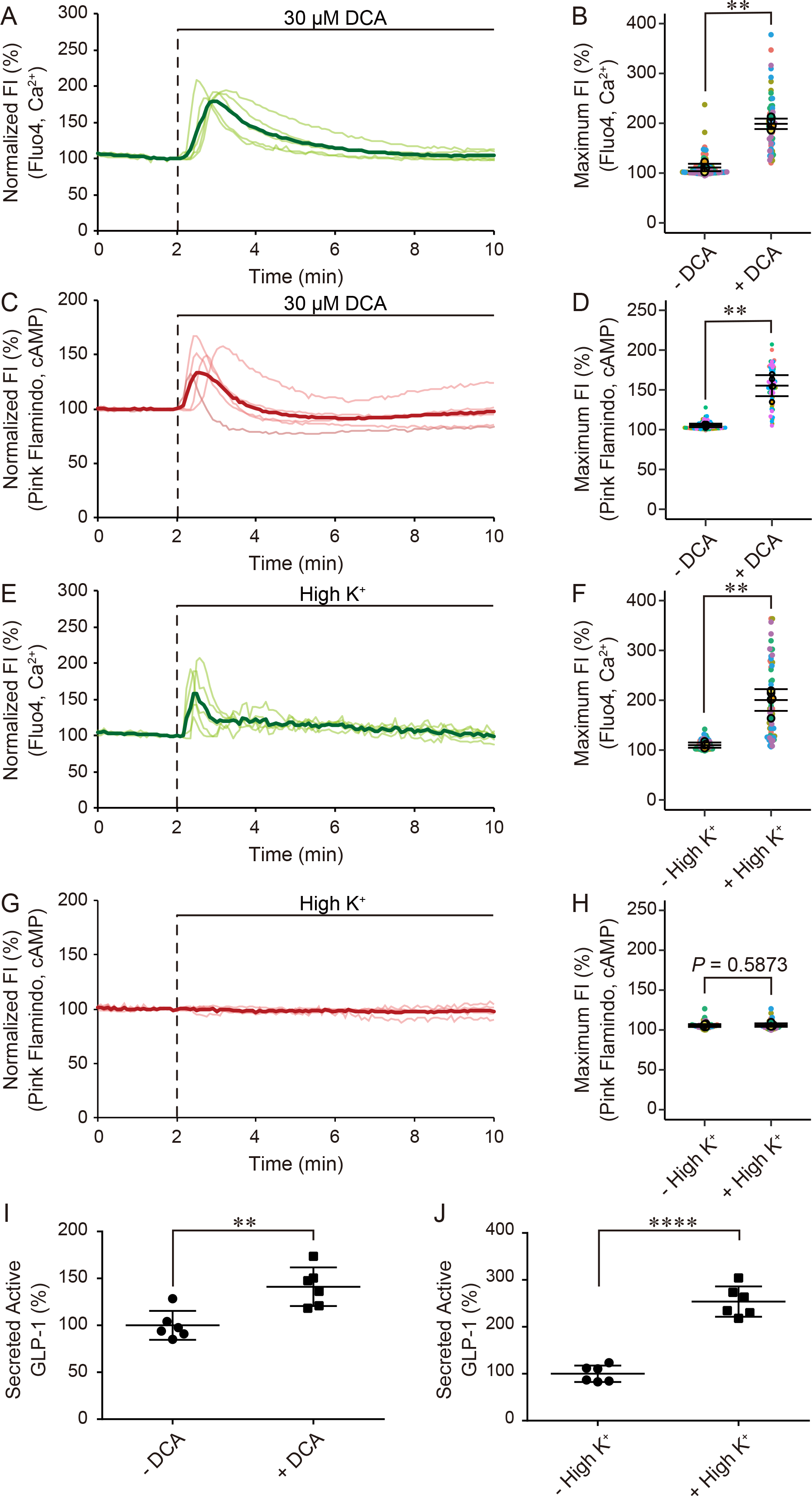
Effect of DCA and high K^+^ on intracellular Ca^2+^ and cAMP dynamics in GLUTag cells. (A) Time courses of fluorescence intensity (FI) of Fluo4 during application of 30 μM DCA. (B) Maximum FI of Fluo4 by application of DCA. (C) Time courses of FI of Pink Flamindo during application of 30 μM DCA. (D) Maximum FI of Pink Flamindo by application of DCA. (E) Time courses of fluorescence intensity (FI) of Fluo4 during application of high K^+^ (15.2 mM KCl). (F) Maximum FI of Fluo4 by application of high K^+^. (G) Time courses of FI of Pink Flamindo during application of high K^+^. (H) Maximum FI of Pink Flamindo by application of high K^+^. (I, J) The amount of secreted active GLP-1 in GLUTag cells after application of 30 μM DCA (I) or high K^+^ (J). Data are shown as means ± SD from six independent experiments. For time courses, Average of normalized FI in single cells are calculated per dish (pale lines). Thick lines represent the average of five or six independent experiments. For super plots, data are shown as means ± SD between five or six experiments. Mann-Whitney U test. ** *p* < 0.01, **** *p* < 0.0001.

We next investigated the signaling pathways derived from DCA and high K^+^. DCA can activate the G protein-coupled bile acid receptor TGR5 which is coupled G_s_ protein (Parker et al., 2012), and also can enter the cells via apical sodium-dependent bile acid transporter (ASBT). We confirmed the expression of *Tgr5* and *Asbt* mRNA in GLUTag cells by RT-PCR (Fig. S1A). Since previous studies have not completely discovered the source of [Ca^2+^]_i_ elevation by DCA (Brighton et al., 2015a), we focused on the ryanodine receptor (RyR) as another putative candidate. When we applied RyR antagonist dantrolene with DCA, FI increase of Fluo4 was significantly suppressed (Fig. 2A, B). In contrast, removal of extracellular Ca^2+^ had little effect on DCA-induced FI increase of Fluo4 (Fig. S1B, C). Inhibition of adenylyl cyclase by 2’,5’-dideoxyadenosine (DDA) significantly suppressed the FI increase of Pink Flamindo (Fig. 2C, D). As for high K^+^, removal of extracellular Ca^2+^ significantly suppressed the FI increase of Fluo4 (Fig. 2E, F). These results suggest that [Ca^2+^]_i_ and [cAMP]_i_ increase by DCA increase derives from activation of RyR and TGR5-mediated G_s_ protein, respectively. [Ca^2+^]_i_ increase by high K^+^ may derive from influx of extracellular Ca^2+^ following membrane depolarization.

**Figure 2.**
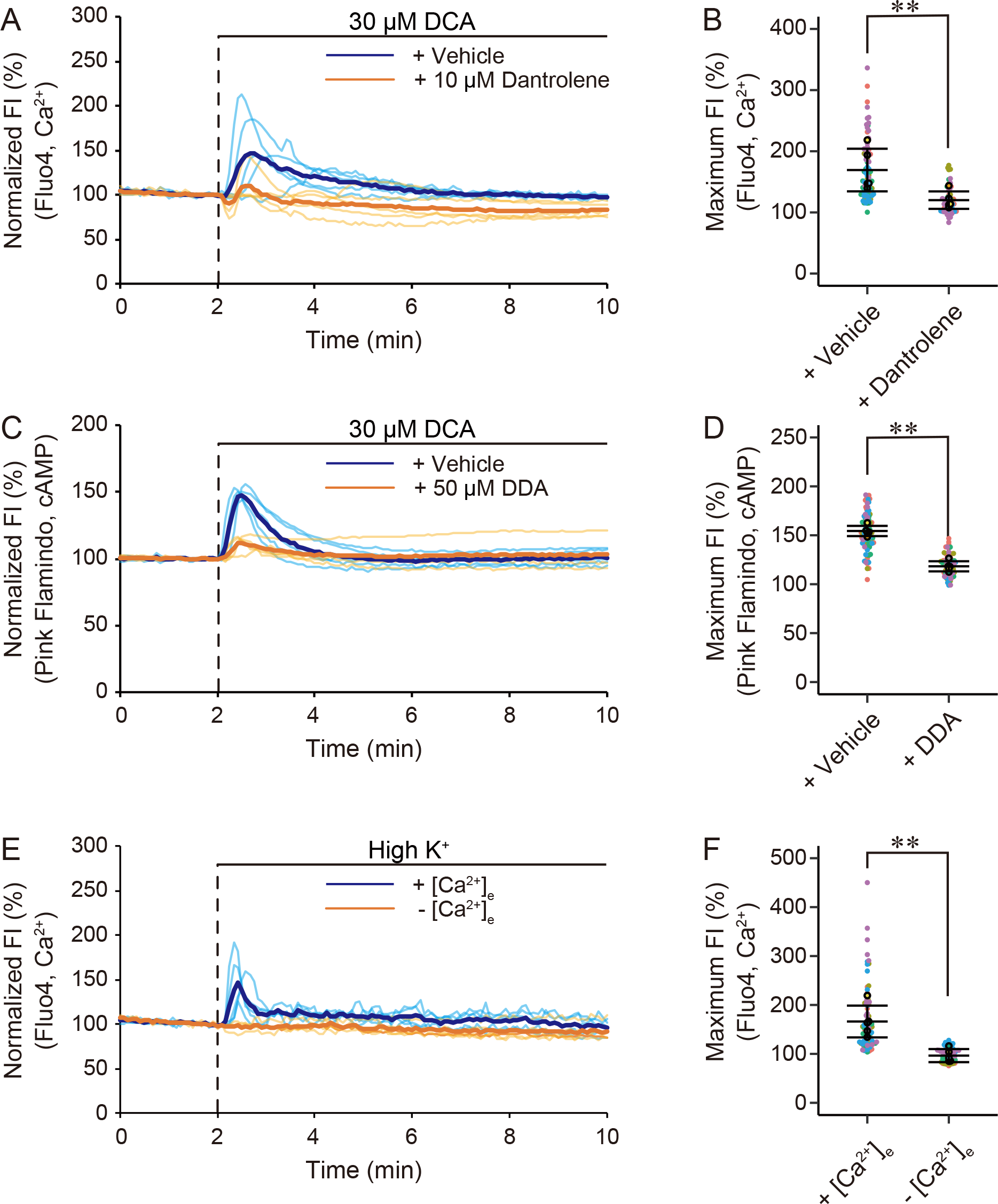
Involvement of ryanodine receptor and G_s_ signaling pathways in DCA-induced [Ca^2+^]_i_ and [cAMP]_i_ increase, and influx of extracellular Ca^2+^ in high K^+^-induced [Ca^2+^]_i_ increase. (A) Time courses of fluorescence intensity (FI) of Fluo4 during co-application of 10 μM dantrolene with 30 μM DCA. (B) Maximum FI of Fluo4 by co-application of dantrolene with DCA. (C) Time courses of FI of Pink Flamindo during co-application of 50 μM DDA with 30 μM DCA. (D) Maximum FI of Pink Flamindo by co-application of DDA with DCA. (E) Time courses of FI of Fluo4 during application of high K^+^ under removal of extracellular Ca^2+^ ([Ca^2+^]_e_). (F) Maximum FI of Fluo4 by high K^+^ under removal of extracellular [Ca^2+^]_e_. Mann-Whitney U test. ** *p* < 0.01.

Previous studies have pointed out [Ca^2+^]_i_ increase by bile acids in pancreatic acinar cells and primary cultured L cells (Brighton et al., 2015b; Voronina et al., 2002), but its molecular mechanisms are not well understood. We propose that DCA is taken up via ASBT and activates RyR to induce Ca^2+^ release from the endoplasmic reticulum, although the precise relationship between DCA and RyR remains unclear. Given that DDA and dantrolene had little effect on FI increase of Fluo4 and Pink Flamindo, respectively (Fig. S1D to G), effect of DCA on [Ca^2+^]_i_ through RyR may be independent of TGR5 activation.

### Exocytotic dynamics of GLP-1 granules show a time-dependent pattern regardless of stimuli

We examined the contribution of [Ca^2+^]_i_ and [cAMP]_i_ increase to GLP-1 exocytosis with total internal reflection fluorescence microscopy (TIRFM). In order to monitor a single exocytosis, we expressed tissue-type plasminogen activator (tPA)-GFP, which well colocalized with endogenous GLP-1 in GLUTag cells (Fig. 3A) (Oya et al., 2013). tPA-GFP shows a relatively slow dynamics in its FI as seen in the previous study (Suzuki et al., 2009). Each exocytotic process can be categorized into three types (Fig. 3B): those predocked with the plasma membrane before stimulation (Old face), docked with the plasma membrane after stimulation and eventually fused (Resting newcomer), rapidly recruited and fused without stable docking to the plasma membrane (Restless newcomer) (Shibasaki et al., 2007). We compared the proportion of these three types at 2 min intervals for 30 min. Exocytotic dynamics showed a biphasic pattern similarly with insulin, where the first and second peak appears within around 8 min and 15 min after stimulation, respectively (Nunemaker et al., 2006; Stožer et al., 2021; Wang et al., 2020). We thus defined the first 8 min from stimulation as “First phase” and thereafter as “Second phase”. Interestingly, we found that Old face and Restless newcomer mainly during the first phase, and Resting newcomer became dominant during the second phase by both DCA and high K^+^ (Fig. 3C, D). Proportion of Resting newcomer significantly increased in the second phase compared to the first phase (Fig. 3E, F), while those of Old face and Restless newcomer tended to decrease (Fig. S2A to D). Although the kinetics of exocytosis frequency appeared different between DCA and High K^+^, proportion of each exocytosis type in total 30 min was not different (Fig. S2E to G). These results suggest that GLP-1 exocytosis immediately after stimulation mainly consists of Old face and Restless newcomer, and shifts to Resting newcomer independent of stimuli and downstream signals.

**Figure 3.**
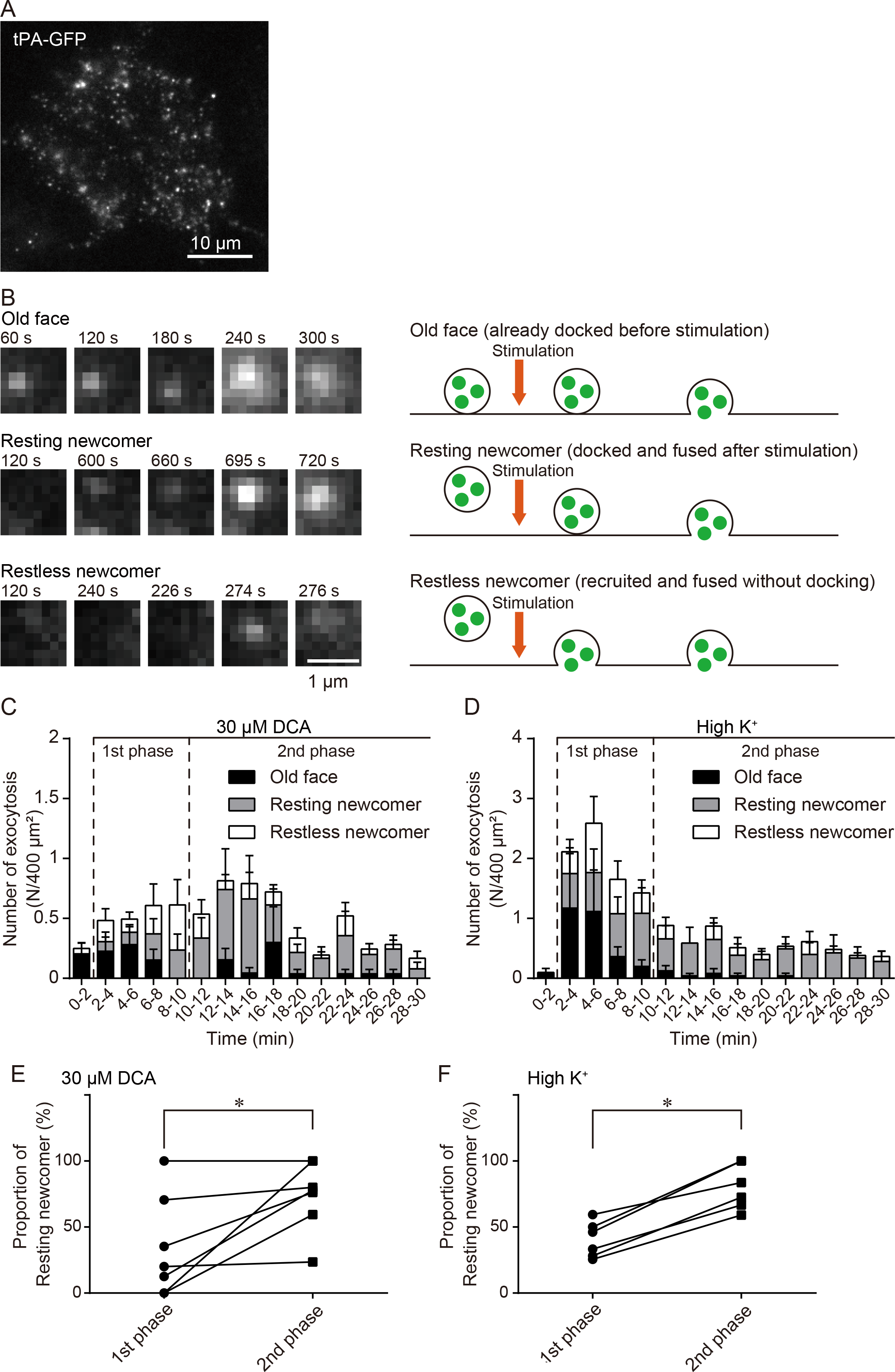
Exocytotic dynamics of GLP-1 from GLUTag cells by DCA and high K^+^. (A) Typical TIRF image of a tPA-GFP-expressing GLUTag cell. (B) Left, typical sequential images of the single granule with Old face, Resting newcomer, and Restless newcomer exocytosis. Stimulation was applied at 120 s. Right, schematic diagrams representing each exocytosis type. (C, D) Histogram of exocytosis observed under TIRFM during application of 30 μM DCA (C) and high K^+^ (D). Data are shown as means ± SEM. N = 13 (C) and 10 (D) cells. (E, F) Proportion of Resting newcomer exocytosis during the first phase (2–10 min) vs second phase (10–30 min) in the cells shown in C and D, respectively. Data are shown as means ± SD. N = 7 (C) and 6 (D) cells. Cells with no exocytosis during the first or second phase were removed from comparison. Wilcoxon matched-pairs signed rank test. * *p* < 0.05.

Previous studies with pancreatic β cells elaborately described the dynamics of insulin exocytosis by different stimuli and diverse genetic backgrounds (Kasai et al., 2008; Mizuno et al., 2016; Shibasaki et al., 2007; Wang et al., 2020). Insulin exocytosis shows a marked difference in the proportion of Old face, Resting newcomer, and Restless newcomer depending on stimuli. For example, high K^+^ induced Old face immediately after stimulation, while glucose induced Restless newcomer for a much longer period (Kasai et al., 2008; Shibasaki et al., 2007). Moreover, elevation of [cAMP]_i_ by 8-Bromo-cAMP in addition to glucose increased the frequency of Restless newcomer (Shibasaki et al., 2007). Since glucose increases not only [Ca^2+^]_i_ but also [cAMP]_i_ through activation of Ca^2+^-sensitive adenylyl cyclases and metabolic production, cAMP seems to enhance Restless newcomer in insulin exocytosis. Regarding GLP-1, however, precise determination of exocytotic dynamics has not been achieved. We found that GLP-1 exocytosis pattern was different from that of insulin and surprisingly independent of stimuli. Because stimulation which only increases [cAMP]_i_ can be an additional approach to gain more insight on the contribution of cAMP, artificial regulation of cAMP production by photoactivated adenylyl cyclase (Stierl et al., 2011), for example, may be a helpful tool to further investigate the role of cAMP with higher resolution.

### Actin polymerization regulates the proportion of exocytosis types

In addition to second messengers, cytoskeletal elements, especially actin network, play pivotal roles in the regulation of exocytosis (Miklavc and Frick, 2020). We thus focused on the involvement of cortical filamentous actin (F-actin) network in GLP-1 exocytosis from GLUTag cells. When we applied 1 μM latrunculin A (Lat A, inhibitor of actin polymerization) to GLUTag cells, cortical F-actin was remarkably disrupted without significant effect on [Ca^2+^]_i_ and [cAMP]_i_ (Fig. S3A to C). We next applied Lat A with DCA and observed exocytotic dynamics with TIRFM. We found that co-application of Lat A lowered the frequency of Old face and Restless newcomer especially during the first phase (Fig. 4A, B). Proportion of Old face in total 30 min was significantly lower, and that of Resting newcomer was significantly higher in the presence of Lat A (Fig. 4C, D). In contrast, proportion of Restless newcomer was not different (Fig. 4E). These results suggest that actin polymerization is essential for facilitation of Old face exocytosis immediately after stimulation, while Restless newcomer may be regulated by other factors.

**Figure 4.**
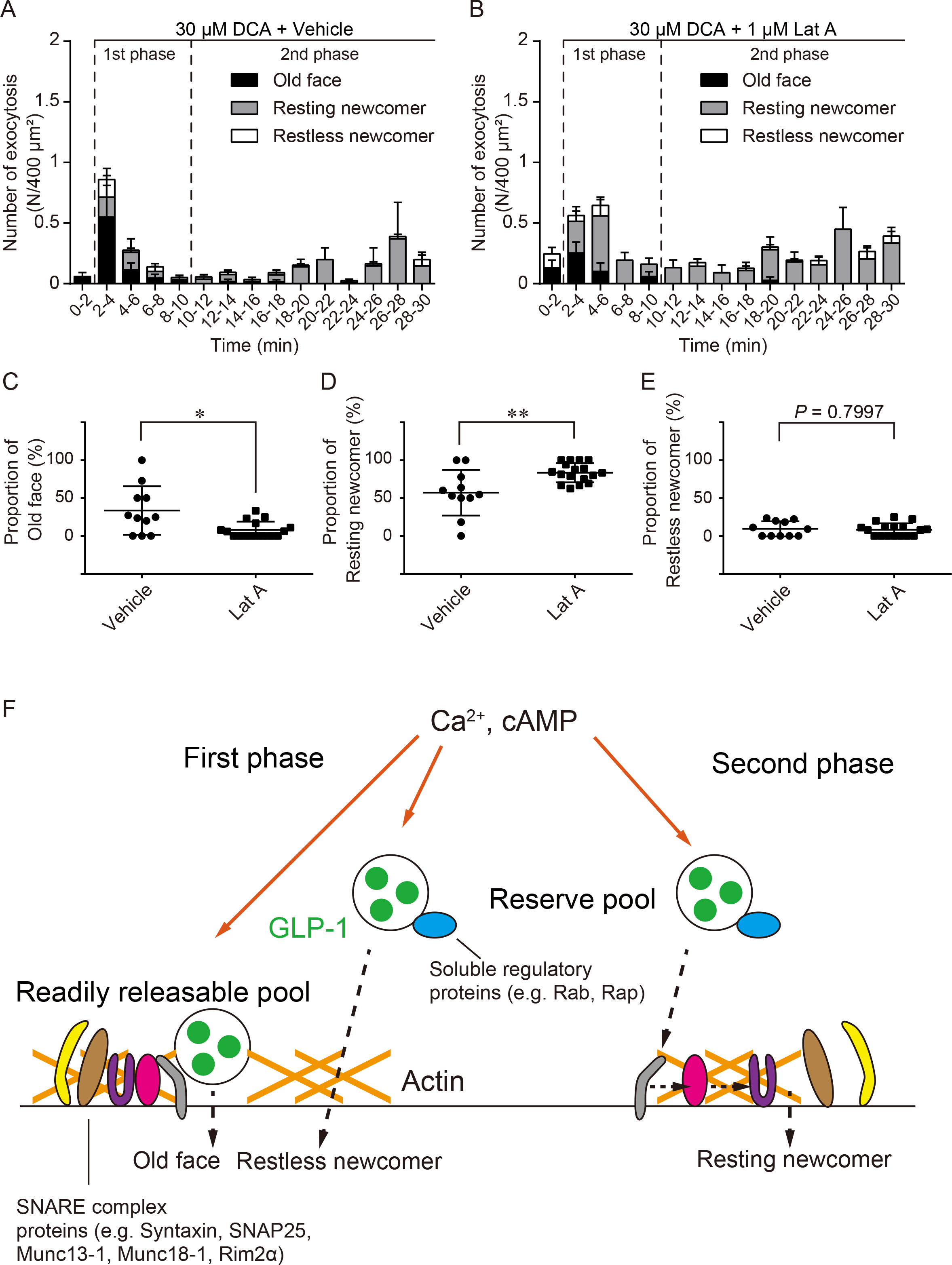
Involvement of F-actin in the regulation of exocytotic dynamics of GLP-1. (A, B) Histogram of exocytosis observed under TIRFM during application of 30 μM DCA plus vehicle (DMSO) (A) and 1 μM Lat A (B). Data are shown as means ± SEM. N = 26 (A) and 18 (B) cells. (C to E) Proportion of Old face (C), Resting newcomer (D), and Restless newcomer (E) exocytosis in the cells shown in A and B. Data are shown as means ± SD. N = 11 (vehicle) and 17 (Lat A) cells. Cells with no exocytosis were removed from comparison. (F) Schematic diagram of a hypothetical model for stimulation-induced GLP-1 exocytosis. In the first phase, cortical F-actin is associated with SNARE complex and facilitates rapid priming and fusion of predocked vesicles, leading to Old face. Restless newcomer may be regulated by other factors than F-actin including soluble regulatory proteins (e.g., Rab, Rap). In the second phase, SNARE complex is dissociated from complex-actin, and newly recruited vesicles together with soluble regulatory proteins are stepwise docked, primed, and fused as Resting newcomer. Ca^2+^ and cAMP are relevant to induction of all exocytosis types in both phases. Mann-Whitney U test. * *p* < 0.05, ** *p* < 0.01.

Role of F-actin in the regulation of exocytosis is controversial: while it acts as a physical barrier between secretory granules and the plasma membrane, it also provides a track for the granules to access the plasma membrane and a scaffold for various regulatory proteins on the fusion cites (Kalwat and Thurmond, 2013; Wang and Thurmond, 2009). In pancreatic β cells, insulin-containing granules are located at two different compartments, the readily releasable pool (RRP) and the reserve pool (RP). Cortical F-actin works as a track for RP-located granules to enter the RRP via interaction with myosin. It has also been reported that the RRP is the main source of Old face during the first phase, and newly recruited granules from the RP accounts for the Resting and Restless newcomer during the second phase (Kasai et al., 2008). In addition, actin and microtubule networks provide “hotspots” for spatially clustered Resting and Restless newcomer of insulin from pancreatic β cell line INS-1 cells (Yuan et al., 2015).

Our results with GLP-1-secreting GLUTag cells shows a difference compared to pancreatic β cells. Although actin-interacting proteins including Rac1 and Cdc42 are required for second phase insulin secretion (Wang et al., 2007), effect of Lat A on the exocytotic dynamics of GLP-1 appears stronger in the first phase. We speculate that the primary role of cortical F-actin in GLUTag cells is to accelerate rapid priming and fusion in the first phase, resulting in the higher proportion of Old face than in the second phase. Priming-promoting proteins such as syntaxins, SNAP25, Munc13-1, Munc18-1, and Rim2α can form a large complex to facilitate priming of adjacent granules, and F-actin can interact with those proteins (Li et al., 2014; Miklavc and Frick, 2020; Toonen et al., 2006; Yasuda et al., 2010). Interestingly, syntaxins and SNAP25 dissociate from F-actin after 5–10 min of glucose stimulation but again associates within 30 min in pancreatic β cell line MIN6 cells (Jewell et al., 2008; Thurmond et al., 2003; Wang and Thurmond, 2009). It is possible that a similar association/dissociation machinery of F-actin and SNARE complex but with different kinetics exists in GLUTag cells. We design a hypothesis for actin-mediated regulation of GLP-1 exocytosis as follows; (i) associated F-actin and SNARE complex provide a scaffold for rapid priming and fusion to predocked granules in the RRP (Old face) in the first phase, (ii) some granules coming from the RP (Restless newcomer) in the first phase may be regulated by soluble regulatory proteins including Rab, Rap, granuphilin (Mizuno et al., 2016; Stožer et al., 2021) rather than F-actin, (iii) after dissociation of SNARE complex from F-actin, granules from RP are docked and primed step-by-step (Resting newcomer) in support of soluble regulatory proteins. Ca^2+^ and cAMP are important for inducing fusion in both phases and all exocytotic types (Fig. 4F). Further investigation into biochemical interaction and multi-color visualization of subcellular localization between the mentioned proteins will enable more knowledge about this model.

## Conclusion

In the present study we investigated the contribution of intracellular Ca^2+^, cAMP, and cortical F-actin to stimulus-induced GLP-1 exocytosis from GLUTag cells. Either increase of [Ca^2+^]_i_ only or increase of both [Ca^2+^]_i_ and [cAMP]_i_ resulted in a similar pattern of exocytosis, where Old face and Restless newcomer are dominant in the first phase and Resting newcomer mainly occurs in the second phase. This time-dependent shift of exocytotic dynamics was considered to be associated with cortical F-actin dynamics. Since we have previously found multiple effects of actin remodeling on GLP-1 exocytosis (Harada et al., 2018, 2017b), there should be a strict machinery promoting/restricting exocytosis through actin network, e.g. cortical F-actin and stress fibers/focal adhesions. In future, understanding the overview of regulatory machanisms in GLP-1 exocytosis under more physiological environments, including primary culture experiments with enterocytes, or direct *in vivo* visualization will provide more breakthroughs.

## Materials and methods

### Chemicals

Deoxycholic acid (DCA) was purchased from Nacalai Tesque (Kyoto, Japan). Dantrolene sodium salt and latrunculin A were purchased from FUJIFILM WAKO Pure Chemical Industries (Osaka, Japan). 2’, 5’-dideoxyadenosine (DDA) was purchased from Sigma-Aldrich (St. Louis, MO, USA).

### Plasmid construction

Green fluorescent protein-tagged tissue-type plasminogen activator (tPA-GFP) and Pink Flamindo plasmids were constructed as described previously (Harada et al., 2017a; Oya et al., 2013).

### Cell culture and plasmid transfection

GLUTag cells (kindly provided by Dr. Daniel Drucker, Lunenfeld-Tanenbaum Research Institute, Canada) were cultured in Dulbecco’s modified Eagle’s medium (Sigma-Aldrich) supplemented with 1 g/L glucose, L-glutamine, sodium pyruvate, 10% (v/v) heat-inactivated fetal bovine serum (Sigma-Aldrich), 100 U/mL penicillin and 100 μg/mL streptomycin (Sigma-Aldrich), at 37°C under 5% CO_2_. For imaging experiments, the cells were trypsinized, and 1×10^5^ cells were plated onto poly-L-lysine (Nacalai tesque)-coated glass coverslips in 35 mm dishes. 2 days after plating, the cells were transfected with 1.5 μg of plasmids using 3 μL of Lipofectamine 2000 Trasnfection Regent (Thermo Fisher Scientific, Waltham, MA, USA) according to the manufacturer’s protocol. The media was exchanged 4 h after transfection and the cells were cultured at 37°C (tPA-GFP) or 32°C (Pink Flamindo) for 2 days until imaging.

### Ca^2+^ and cAMP imaging

For Ca^2+^ imaging, GLUTag cells plated on 35 mm dishes for 2 days were washed twice, and loaded with 250 nM of Fluo 4-AM (Dojindo, Kumamoto, Japan) in modified Ringer Buffer (RB: 140 mM NaCl, 3.6 mM KCl, 0.5mM NaH_2_PO_4_, 0.5mM MgSO_4_, 1.5 mM CaCl_2_, 10 mM HEPES, 2 mM NaHCO_3_) containing 5 mM glucose. After incubation for 20 minutes at 37°C under 5% CO_2_, the cells were washed twice, added with RB containing 0.1 mM glucose. For cAMP imaging, GLUTag cells transfected with Pink Flamindo was simply washed wtice and added with RB containing 0.1 mM glucose. Imaging was performed using an inverted microscope (IX-71, Olympus, Tokyo, Japan) equipped with an oil-immersion objective lens (UApo/340, 40×, NA = 1.35, Olympus), a stage heated at 37°C, and an EM-CCD camera (Evolve, Photometrics, Tucson, AZ, US) whose exposure was controlled by MetaMorph software (Molecular Devices, Sunnyvale, CA, USA). Images were acquired using a xenon lamp, 460–495 nm excitation filter, 505 nm dichroic mirror and 510–550 nm emission filter (U-MWIBA2, Olympus) every 5 s for 10 min. Stimuli were applied at 120 s from the beginning of image acquisition. DCA was dissolved in dimethyl sulfoxide (DMSO) and applied at the final concentration of 30 μM. High K^+^ solution consisted of 128.4 mM NaCl, 15.2 mM KCl, 0.5mM NaH_2_PO_4_, 0.5mM MgSO_4_, 1.5 mM CaCl_2_, 10 mM HEPES, 2 mM NaHCO_3_, 0.1 mM glucose.

For inhibition experiments, DDA and latrunculin A were co-applied DCA simultaneously, while dantrolene was applied 20 min before the beginning of image acquisition. Removal of extracellular Ca^2+^ was performed with Ca^2+^-free RB (140 mM NaCl, 3.6 mM KCl, 0.5mM NaH_2_PO_4_, 0.5mM MgSO_4_, 10 mM HEPES, 2 mM NaHCO_3_, 2.5 mM EGTA, 0.1 mM glucose).

### Enzyme-linked immunosorbent assay

GLUTag cells were plated in 24-well plates at 1×10^5^ cells per well. Two days after plating, cells were washed twice with RB containing 5 mM glucose. Then, either control solution (0.03% v/v DMSO for DCA, no additives for high K^+^), DCA, or high K^+^ in RB containing 0.1 mM glucose were applied to the cells and incubated for 1 h at 37°C under 5% CO_2_. After centrifugation at 1,000 g for 10 min at 4 °C, supernatant was used for analysis with GLP-1, Active form Assay Kit (Immuno-Biological Laboratories, Gunma, Japan) and microplate reader (Varioskan LUX, Thermo Fisher Scientific).

### RNA isolation and RT-PCR

RNA isolation, reverse transcription, and PCR were performed as described previously (Harada et al., 2017b). For amplification of glyceraldehyde 3-phosphate dehydrogenase (*Gapdh*, NM_001289726.1), the forward primer 5′-GGAAGGGCTCATGACCACAG-3′ and the reverse primer 5′-ACCAGTGGATGCAGGGATGA-3′ were used. For G protein-coupled bile acid receptor 1 (*Tgr5*, NM_174985.1), the forward primer 5′-GCTACATGGCAGTGTTGCAG-3′ and the reverse primer 5′-CTGGGAAGACAGCTTGGGAG-3′ were used. For apical sodium-dependent bile acid transporter (*Asbt*, NM_011388.3), the forward primer 5′-GCGACATGGACCTCAGTGTT-3′ and the reverse primer 5′-GTTCCCGAGTCAACCCACAT-3′ were used.

### Total internal reflection fluorescence microscopy (TIRFM)

GLUTag cells transfected with tPA-GFP were washed and imaged as described above. Imaging was performed using an inverted microscope (ECLIPSE Ti-E, Nikon, Tokyo, Japan). We used a high numerical aperture objective lens (CFI Apochromat TIRF, 100×, NA=1.49, Nikon), and incident light for total internal reflection illumination was introduced from the objective lens through a single-mode optical fiber and two illumination lenses (TI-TIRF, Nikon). An optically pumped semiconductor 488-nm laser (Sapphire 488LP, 30 mW, Coherent, Santa Clara, Canada) was used through a filter set (465–495 nm excitation filter, 505 nm dichroic mirror, and 515–555 nm emission filter, Nikon) and an electromagnetically driven shutter (TI-TIRF, Nikon), and the shutter was opened synchronously with an EM-CCD camera (iXon, Andor, Belfast, UK), whose exposure was controlled by MetaMorph software. Images were acquired every 500 ms for 30 min.

### Phalloidin staining and confocal imaging

GLUTag cells plated on 35 mm dishes for 2 days were washed twice, applied with control solution (0.1% v/v DMSO) or 1 μM of latrunculin A in RB containing 0.1 mM glucose, and incubated for 10 min at 37°C under 5% CO_2_. Then the cells were washed twice with phosphate-buffered saline (PBS) and fixed with 2% paraformaldehyde in PBS for 20 min at room temperature. After washing 3 times with PBS and permeabilization with 0.3% Triton X-100 in PBS for 2 min, the cells were treated with 0.1% v/v fluorescence-labeled phalloidin (Alexa Fluor^®^ 568 Phalloidin, Thermo Fisher Scientific) in PBS for 30 min at room temperature in dark. Confocal images were acquired to investigate their localization using a laser confocal microscope (C2◻+, Nikon, Tokyo, Japan) equipped with an oil immersion◻100× objective lens (Plan Apo VC,◻100×, NA◻=◻1.40, Nikon) and 561 nm laser (Sapphire, Coherent, Santa Clara, CA, USA).

### Imaging data analysis

Acquired images for imaging experiments were analyzed using ImageJ (National Institutes of Health, Bethesda, MD, USA) and MetaMorph software. XY drifts of images were corrected using “Stackreg” plugin in ImageJ if necessary. For Ca^2+^ and cAMP imaging, basal fluorescence intensity, normalized as 100%, was calculated as an average of fluorescence intensity (FI) during 30 s immediately before stimulation, and maximum FI after stimulation was calculated. Then, average of maximum FIs in each trial was calculated and used for statistical comparisons. Individual data are represented as super plots using RStudio. Each large dot represents the average of single cells (small dots) with the same color. For TIRFM with tPA-GFP, we focused on changes in FI of single vesicles to distinguish between exocytosis types. Individual vesicles were selected manually, and the average FI of a single vesicle in a 1.3×1.3 μm square over the vesicle center was measured. Exocytosis was defined as: (i) FI is elevated by ≥ 30% (priming and fusion with the plasma membrane) within 20 s from the time when a single vesicle is detected in the cell area (recruit and docking), (ii) FI declines over time after reaching a peak (release of vesicle contents), and (iii) the vesicle is tethered within a 1.3×1.3 μm square during FI elevation and subsequent decline. We then classified each exocytosis into three categories. If a vesicle which is already docked with the plasma membrane before stimulation and fused, it is defined as “Old face”. If a vesicle is docked after stimulation for 2 frames (1 s) or more and subsequently fused, it is defined as “Resting newcomer”. If a vesicle is rapidly fused within a single frame after appearance, it is defined as “Restless newcomer”. We recognized few Old face and Restless newcomer events before stimulation. The number of each type of exocytosis (N /400 μm^2^) was calculated for each cell.

For statistical analysis, data are shown as means ± standard deviation or standard error of mean. Means were compared by Welch’s *t* test or paired *t* test if the normal distribution of groups are guaranteed by D’Agostino & pearson omnibus test, otherwise Wilcoxon single rank test or Wilcoxon matched pairs singled rank test using GraphPad Prism 6 software (GraphPad software, La Jolla, CA, USA).

## Abbreviations

GLP-1: glucagon-like peptide-1
GIP: glucose-dependent insulinotropic polypeptide
DCA: deoxycholic acid
SNARE: soluble NSF attachment protein receptor
FI: fluorescence intensity
ASBT: apical sodium-dependent bile acid transporter
Lat A: latrunculin A
RRP: readily releasable pool
RP: reserve pool

## Acknowledgments

We thank Dr. Daniel Drucker for kindly providing GLUTag cells and Dr. Yongdeng Zhang for helpful advice on exocytosis analysis.

