## Supplementary figures for "Exocytotic dynamics of glucagon-like peptide-1 from enteroendocrine L cell line is regulated by actin polymerization"

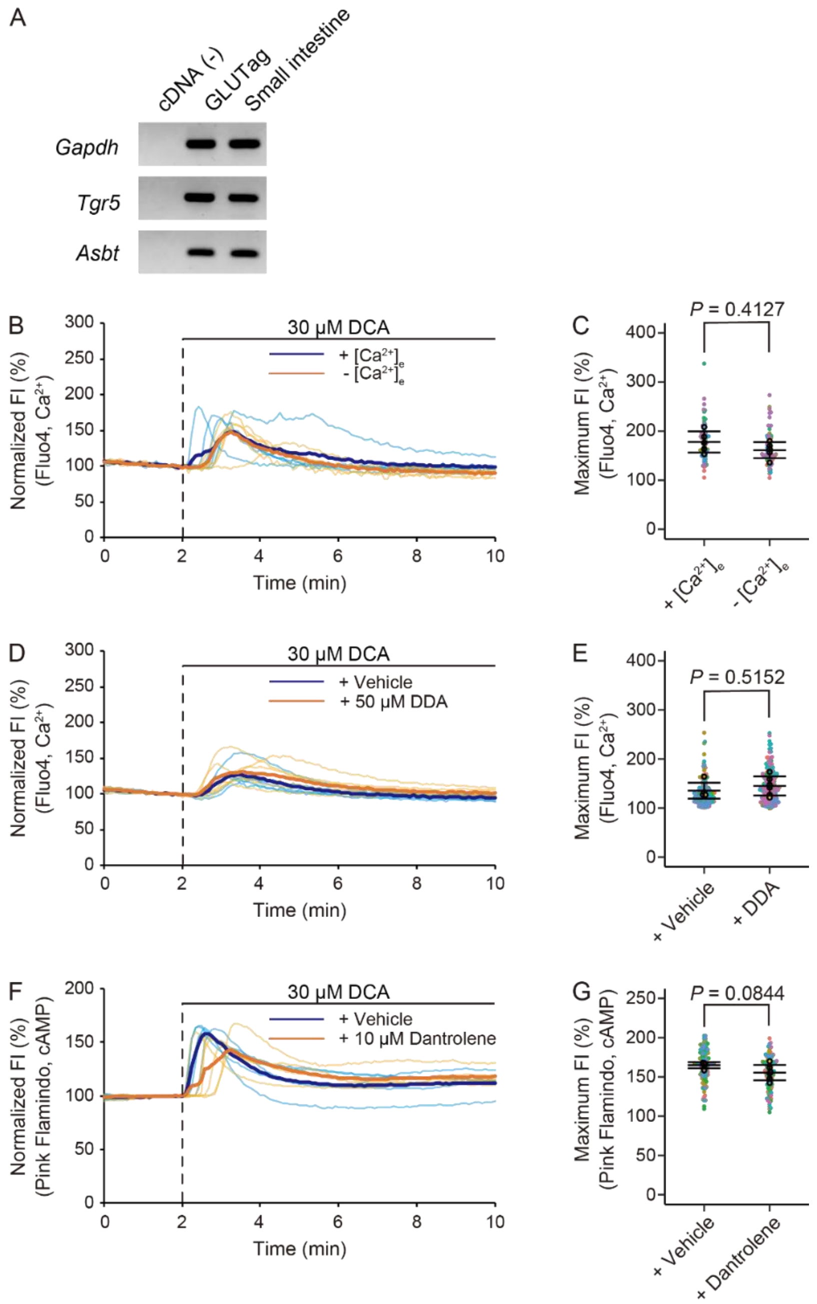


**Figure S1. Expression of *Tgr5* and *Asbt* in GLUTag cells, and additional inhibition experiments on DCA and high $K^+$ application.**

(A) RT-PCR analysis of mRNA expression of *Tgr5* and *Asbt* in GLUTag cells. N = 3 independent trials. (B) Time courses of fluorescence intensity (FI) of Fluo4 during application of 30 µM DCA under removal of extracellular $Ca^{2+}$ ($[Ca^{2+}]_e$). (C) Maximum FI calculated from the FI of Fluo4 by DCA under removal of extracellular $[Ca^{2+}]_e$. (D) Time courses of FI of Fluo4 during co-application of 50 µM DDA with 30 µM DCA. (E) Maximum FI calculated from the FI of Fluo4 by co-application of DDA with DCA. (F) Time courses of FI of Pink Flamindo during co-application of 10 µM dantrolene with 30 µM DCA. (G) Maximum FI calculated from the FI of Pink Flamindo by co-application of dantrolene with DCA.

Time courses and super plots are illustrated in the same manner as Main Figure 1. Mann-Whitney U test.

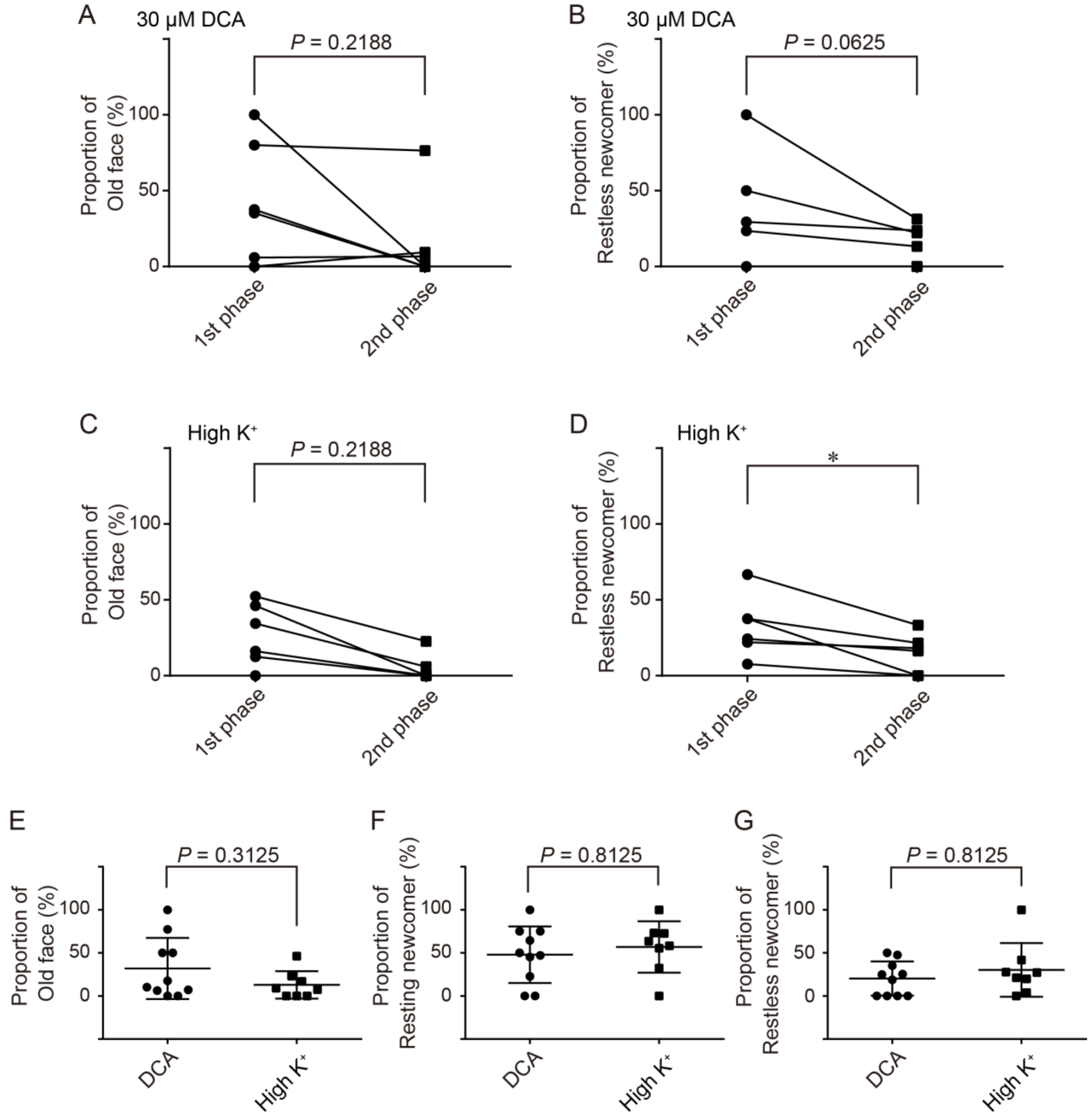


**Figure S2. Additional information about the exocytotic dynamics of GLP-1 from GLUTag cells by DCA and high K$^+$.**

(A to D) Proportion of Old face (A, C) and Restless newcomer (B, D) exocytosis during the first phase (2–10 min) vs second phase (10–30 min) in the cells shown in Main Figure 3C and D. Data are shown as means ± SD. N = 7 (C) and 6 (D) cells. It is noted that cells with no exocytosis during the first or second phase were removed from comparison. (E to G) Proportion of Old face (E), Resting newcomer (F), and Restless newcomer (G) exocytosis in the cells shown in Main Figure 3C and D. Data are shown as means ± SD. N = 10 (DCA) and 8 (high K$^+$) cells. It is noted that cells with no exocytosis were removed from comparison. Wilcoxon matched-pairs signed rank test (A to D) and Mann-Whitney U test (E to G). * $p < 0.05$.

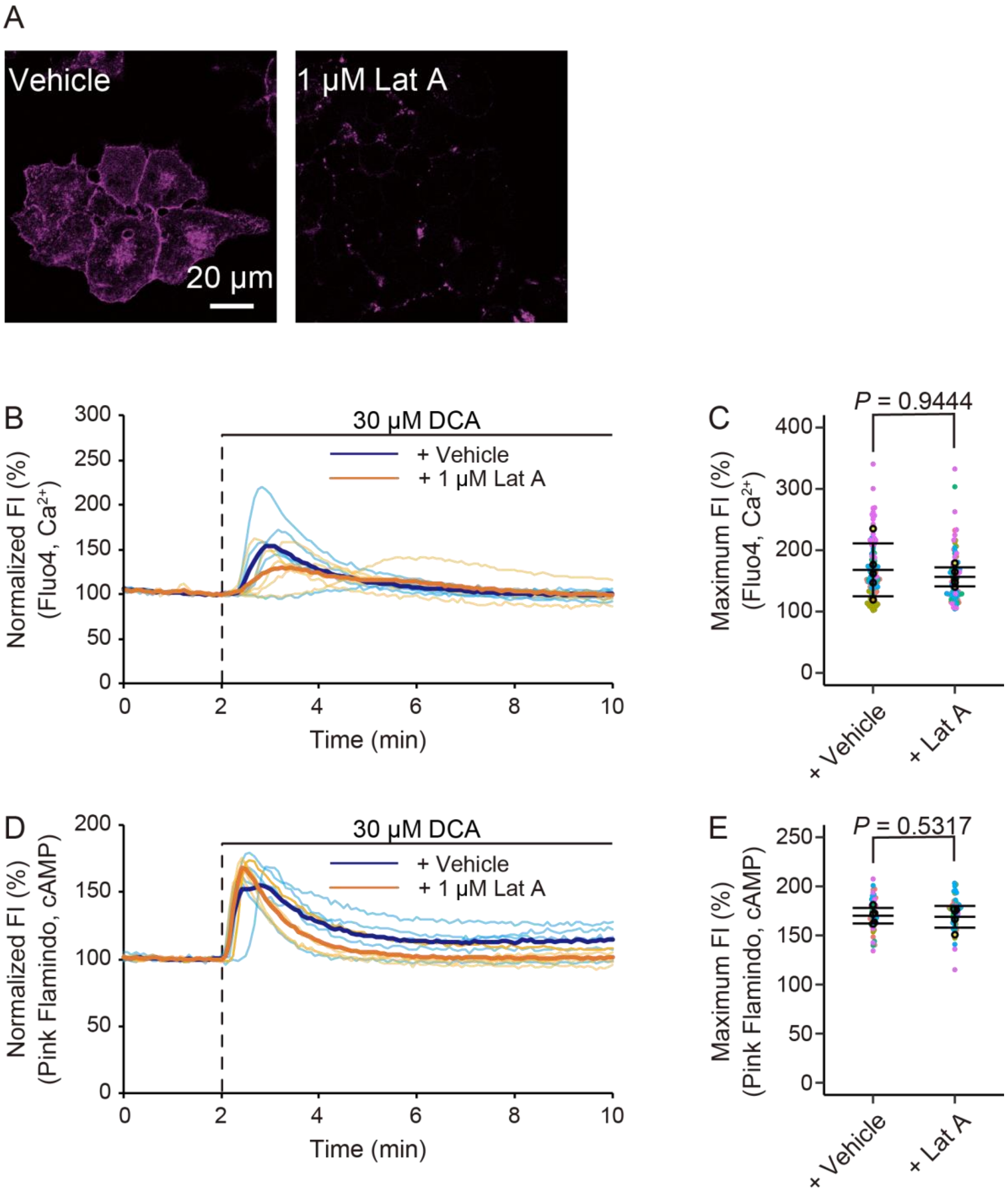


**Figure S3. Effect of latrunculin A (Lat A) on cortical F-actin and DCA-induced $[Ca^{2+}]_i$ and $[cAMP]_i$ increase.**

(A) Typical confocal images of phalloidin-stained GLUTag cells in the presence or absence of 1 μM Lat A. (B) Time courses of fluorescence intensity (FI) of Fluo4 during co-application of 1 μM Lat A with 30 μM DCA. (C) Maximum FI calculated from the FI of Fluo4 by co-application of Lat A with DCA. (C) Time courses of FI of Pink Flamindo during co-application of 1 μM Lat A with 30 μM DCA. (D) Maximum FI calculated from the FI of Pink Flamindo by co-application of Lat A with DCA. Time courses and super plots are illustrated in the same manner as Main Figure 1. Mann-Whitney U test.
